# Intracellular localization and gene expression analysis provide new insights on LEA proteins’ diversity in anhydrobiotic midge

**DOI:** 10.1101/825133

**Authors:** Sabina A. Kondratyeva, Alexander A. Nesmelov, Alexander V. Cherkasov, Yugo Miyata, Shoko Tokumoto, Takahiro Kikawada, Oleg A. Gusev, Elena I. Shagimardanova

## Abstract

Anhydrobiosis, an adaptive ability to withstand complete desiccation, is in insects limited to a single species: the nonbiting midge *Polypedilum vanderplanki* (the sleeping chironomid). Evolution of anhydrobiosis in a single representative of a large genus is associated with drastic changes in genome structure, including the emergence of new multimember gene families directly involved in desiccation tolerance. Among them, Late Embryogenesis Abundant (LEA) proteins, which protect other proteins from aggregation caused by desiccation, are believed to originate via horizontal gene transfer from a bacterial donor. To obtain new insights on the biological background of the expanded 27-member LEA protein group in *P. vanderplanki*, we investigated the expression of corresponding genes in a *P. vanderplanki*-derived cell line, capable of anhydrobiosis, in a normal state and during induction of desiccation tolerance. We found that all LEA proteins genes identified in *P. vanderplanki*’s genome except *PvLea16* and *PvLea17* are also expressed in Pv11 cells. Their expression was elevated in response to anhydrobiosis-inducing trehalose treatment. Expression patterns of *PvLea* genes were well preserved in Pv11 cells in comparison to *P. vanderplanki*’s larvae both in the control group and during the anhydrobiosis cycle. We also investigated localization of LEA proteins in Pv11 cells and Sf9 cells and found a different level of conservation in intracellular localization of the protein expressed in mammalian and insect cells.

## Introduction

One of the most intriguing adaptations to extreme environments is the phenomenon of anhydrobiosis—the ability to withstand complete desiccation in a nonmetabolic state (see review in (1)—which allows living organisms to revive after complete desiccation. Organisms capable of entering anhydrobiosis are found across different taxa, and the larvae of the chironomid *Polypedilum vanderplanki* (Insecta, Diptera) are one of the most complex of these from an evolutionary point of view, as well as being a unique case of emergence of anhydrobiosis in a single species of eukaryotes (2).

For the larvae, entering the anhydrobiotic state takes 48 hours and is mediated by several key events including replacement of water with trehalose and vitrification, as well as accumulation of protective biomolecules, including heat shock proteins, antioxidants and enzymes, aquaporins, and Late Embryogenesis Abundant (LEA) proteins (2,3). The latter are highly hydrophilic proteins, well known for their protective effect against water deficits, and they are found in anhydrobiotic organisms in both the plant and animal kingdoms (2).

Previously, we identified 27 genes of LEA proteins (*PvLea*) in the *P. vanderplanki* genome, all of which are encoding members of LEA group 3. Many of these genes become induced during desiccation in larvae (4). LEA proteins are believed to be one of the main participants in anhydrobiosis mechanisms, but the origin of PvLEA-coding gene diversity, and the function of this diversity in a single species, is still unknown.

To gain further understanding of PvLEA protein functions in *P. vanderplanki*, we performed an RNA-seq analysis of the Pv11 cell line established from embryonic stem cells of *P. vanderplanki* and capable of entering anhydrobiosis (5,6). We also investigated patterns of intracellular localization in the Sf9 cell line. We found incomplete preservation of PvLEA proteins’ localization in comparison to our previous data on exogenous expression of LEA proteins in mammalian cells, which should be taken into account in the application of *P. vanderplanki*-derived PvLEA proteins in dry preservation technologies.

## Materials and methods

### Cell lines

Pv11 cells, originally isolated from *P. vanderplanki* egg masses, were cultivated in accordance with previously published protocols (5). Briefly, we cultivated the cells in the IPL-41 medium (Gibco, USA), supplemented with 10% fetal bovine serum (Hyclone, USA) and 2.6 g/l tryptose phosphate broth. We obtained insect Sf9 cells from Merck and cultivated them in the Sf900 medium (Gibco, USA) without supplements. We maintained both Pv11 and Sf9 cultures in nonhumidified incubators at 25 °C and 28 °C, respectively.

### Expression vectors

We used *PvLea* genes previously cloned from *P. vanderplanki* larvae into *PvLea*-AcGFP1 fused genes (7), and we excised these fused genes from pcDNA5/FRT-PvLeaX-AcGFP1 (X = 1-27) vectors (7) with BamHI and XhoI restriction enzymes and reinserted them into the pP121K vector using the Ligation-Convenience Kit (Nippon gene, Japan). Vector pP121K has previously been obtained through replacement of the PvGapdh-promoter region of pPGK-AcGFP1 vector (8) with the 121 promoter, isolated from *P. vanderplanki* (9). The resulting vectors were sequenced to confirm their identity against previously published versions (4).

### Transfection and protein expression

We transfected the Pv11 cells using Nepa 21 Super Electroporator (NepaGene, Japan) with the following settings: 6 poring pulses at 250 V for 4 ms with 40% voltage decay and positive voltage polarity, followed by 5 transfer pulses at 20 V for 50 ms with 40% voltage decay and alternating polarity (8). We transfected the Sf9 cells using the Escort IV reagent (Sigma, USA) in accordance with manufacturer recommendations. We cultivated both Pv11 and Sf9 cells for 24 after transfection at standard conditions for protein expression.

### Visualization of PvLEA protein localization

We stained Pv11 cells expressing AcGFP1 with CellVue Claret Far Red (Sigma, USA) for cell membrane labeling, staining DNA in both Pv11 and Sf9 with Hoechst 33258 (Sigma, USA). In the case of the PvLEA3 and PvLEA22 proteins, which are localized in the endoplasmic reticulum (ER) or Golgi apparatus, we stained these organelles using the CytoPainter Staining Kit (Abcam, USA). We investigated expression and localization of fused PvLEA-AcGFP1 proteins in cell cultures using a laser confocal microscope LSM 780 (Zeiss, Germany) in the Interdisciplinary Center for Analytical Microscopy of Kazan Federal University.

### RNA-seq

We subjected the Pv11 cells to the established procedure of anhydrobiosis induction via treatment by 600 mM trehalose (Sigma, USA) in water, with the addition of 10% of complete IPL-41 medium for 48 hours (5). We then dried the cells for 10 days in a desiccator, rehydrated them, and cultivated them normally. We took cell samples in the control group, after 24 and 48 hours of trehalose treatment, and after 24 hours postrehydration. RNA was isolated with TRIzol reagent (ThermoScientific, USA).

We quantified total RNA using a Qubit 2.0 fluorometer and estimated RNA quality using Bioanalyzer 2100 (Agilent, USA). We constructed libraries from 300 ng of RNA using NEB Next Ultra RNA kit (New England Biolabs, USA) and isolated Poly(A) mRNA with 24-dT oligonucleotides by means of the NEB Next Poly(A) mRNA Magnetic Isolation Module and fragmented them. After reverse transcription with 6-nucleotide random primers and second-strand cDNA synthesis, we ligated the adapters and amplified the libraries using a universal primer (the same for all libraries) and index primers. Then, we cleaned the libraries using AMPure XP magnetic beads (Beckman Coulter, USA) and quantified them using the Qubit 3.0 fluorometer. We checked the distribution of fragment length using Bioanalyzer 210 and sequenced the obtained libraries on Illumina HiSeq 2500, with 50 bp single-end sequencing.

We downloaded the RNA-seq dataset for *P. vanderplanki*’s larvae, sequences of genome (version 0.9), Anhydrobiosis-Related gene Island (ARID) regions, and respective gene annotations (version 0.91) from http://bertone.nises-f.affrc.go.jp/midgebase/ and merged the gene annotations and genomic sequences. We mapped RNA-seq reads for larvae and PV11 using hisat2 version 2-2.1.0 (10) onto the combined genomic sequences of the ARID regions and the whole genome of *P. vanderplanki*, and we sorted the resulting sam files using samtools 1.9-52 (11), obtaining corresponding counts using HTSeq 0.5.4p3 (12).

We obtained gene counts using merged gene annotation of ARID regions and whole-genome annotation. The use of ARID regions annotation ensures higher accuracy of *PvLea* expression analysis because AUGUSTUS-predicted whole gene annotation lacks 6 out of 27 *PvLea* genes, expression of which was previously confirmed by qPCR (7). *PvLea* annotation in ARID regions corresponds to *PvLea* sequences verified with cDNA cloning (7). To avoid ambiguous read mapping, we discarded genes from whole-genome annotation that matched ARID gene sequences from the resulting merged annotation. Genes were considered as matching when they had identity > 95% and e-value < 10^−60^ in BLASTn (13) to corresponding sequences of ARID-annotated genes. We considered genes as expressed when they had at least 10 counts in 3 or more samples. We obtained the RPKM values of gene expression using edgeR 3.26.8 (14).

## Results

### All known *PvLea* genes except *PvLea16* and *PvLea17* are expressed in Pv11 cells

To analyze gene expression patterns, we produced an RNA-seq dataset for Pv11 cells in the control group and after 48 hours of trehalose treatment. This treatment is needed to induce anhydrobiosis in Pv11 cells. We found that all 27 *PvLea* genes previously identified in the *P. vanderplanki* genome and expressed in its larvae were expressed in Pv11 cells, excepting *PvLea16* and *PvLea17*. Patterns of *PvLea* genes’ expression were similar in Pv11 cells and larvae (S1 Fig). Importantly, the *PvLea* expression pattern was well preserved between Pv11 cells and larvae on the 24 hours time point of anhydrobiosis induction (Fig. 1), which was shown to be the point of highest expression for many PvLEA proteins in *P. vanderplanki* larvae (7), reflecting the high demand for *P. vanderplanki* in these proteins at this point.

**Fig. 1.**
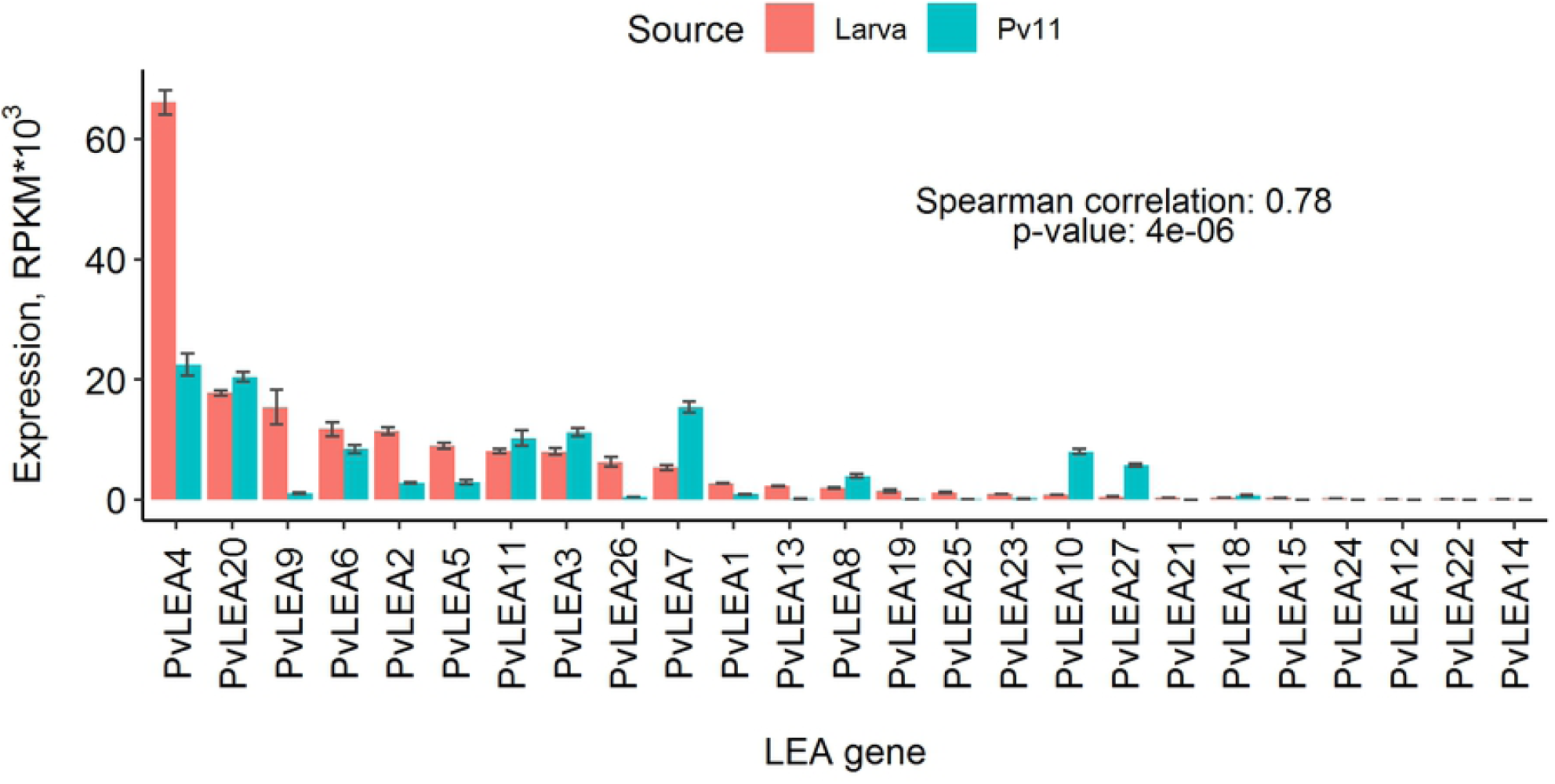
*PvLea* expression in *P. vanderplanki* larvae and Pv11 cells at 24 hours of anhydrobiosis induction. The height of the bars depicts mean expression for replicates; error bars show corresponding standard deviations. The colors of the bars indicate data for larvae (red) or Pv11 cells (blue-green). Genes are ordered in accordance with expression in *P. vanderplanki* larvae, and their IDs are placed below the plot. The text indicates the Spearman correlation values of mean gene expression values between Pv11 cells and *P. vanderplanki* larvae and corresponding *p*-value.

For the *PvLea7, PvLea10, PvLea20*, and *PvLea27* genes, expression was substantially higher in Pv11 cells than in larvae (S1 Fig.). Another remarkable difference in *PvLea* expression between larvae and Pv11 cells was the relatively low expression of *PvLea4* at 24 and 48 hours of anhydrobiosis induction (S1 Fig.).

### PvLea genes become induced in Pv11 cells on a course of anhydrobiosis cycle

We compared expression of PvLEA-expressing proteins at different stages of anhydrobiosis induction. In the case of Pv11, similarly to *P. vanderplanki*’s larvae, *PvLea* genes were induced in response to anhydrobiosis induction in Pv11 cells (Fig. 2). For almost all *PvLea* genes, we observed the highest expression during the trehalose treatment. The *PvLea4* gene had the highest expression levels among all *PvLea* genes.

**Fig. 2.**
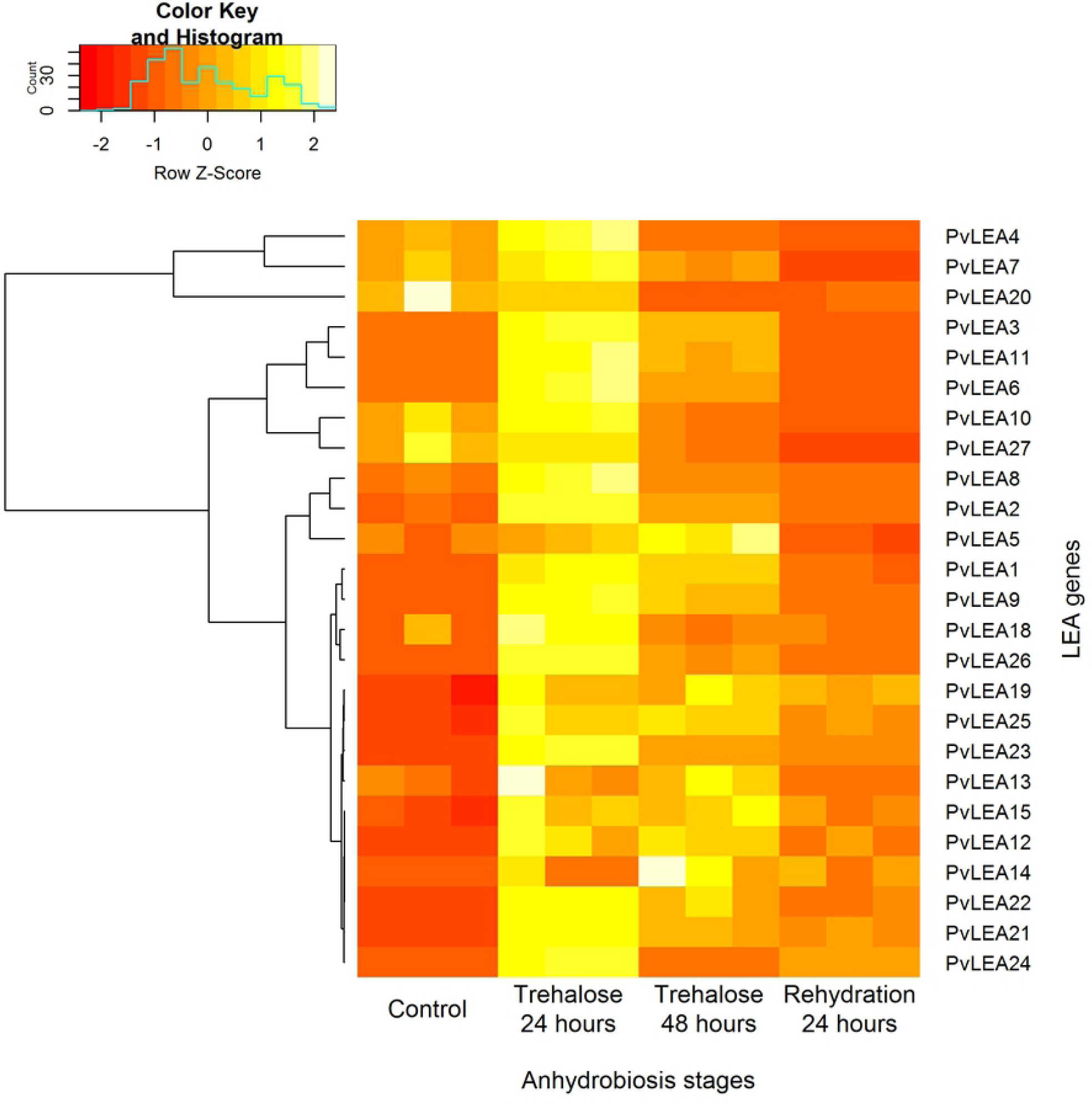
Heat map of *PvLea* genes expression in Pv11 cells. Experimental conditions are indicated at the bottom. We rescaled the expression values for each gene. The resulting values of relative expression for each gene are depicted as rows, with gene names indicated on the right. Color coding of relative expression is on the top of the image, with the brightest and darkest colors reflecting highest and lowest expression for a given gene, respectively. The brackets on the left show the clusters identified based on similarity of relative expression profiles.

We estimated whether some properties of amino acid sequences of PvLEA proteins were correlated with the expression of corresponding *PvLea* genes. For each experimental condition, we computed a Spearman correlation between gene expression and different characteristics of corresponding proteins, including hydropathy (GRAVY, Grand average of hydropathy index), number of LEA_4 (Pfam: PF02987) motifs, and protein disorder degree (FoldIndex) (7). Of these, the number of LEA_4 motifs had the highest correlation with the expression of the corresponding gene (*r* > 0.44, *p*-value 10^−6^ and less) (S1 Table, S2 Fig). High expression of PvLEA4, PvLEA7, and PvLEA20 genes encoding highly hydrophilic proteins with low GRAVY index was also associated with some degree of negative correlation of these characteristics with expression (S1 Table, S2 Fig).

### Subcellular localization of PvLEA proteins in Pv11 and Sf9 cells

To investigate localization of PvLEA proteins in *P. vanderplanki*’s Pv11 cells, we transfected Pv11 cells with 27 vectors containing PvLEA(1-27)-AcGFP1 chimers.

To investigate the intracellular localization of PvLEA proteins, we performed transfection of Pv11 cells with *PvLea*(x)-AcGFP1 genetic vectors (where x = 1-27). We placed these chimers under the control of the 121 promoter, ensuring a high level of gene expression in *P. vanderplanki* cells (9,15).

We targeted all PvLEA proteins according to one of four localization types: whole cell, cytoplasm excluding the nucleus, ER/Golgi, or cellular membrane. Representative images of these localization types are given in Fig. 3. The majority of PvLEA-AcGFP1 fusion proteins, including PvLEA2, PvLEA4, PvLEA9-PvLEA20, PvLEA26, and PvLEA27, in Pv11 cells were distributed across the entire cell, lacking specific localization (Table 1, Fig. 1, and S2 Fig.). Most remaining proteins, including PvLEA5, PvLEA6, PvLEA7, PvLEA8, PvLEA21, PvLEA23, PvLEA24, and PvLEA25, were located in the cytosol of Pv11 cells, clearly excluding the nucleus. Localization of the aforementioned proteins in Sf9 cells was similar, except for the PvLEA7, PvLEA8, PvLEA21, and PvLEA23 proteins, which were not excluded from the nucleus in Sf9 cells (Table 1, Fig. 1, and S2 Fig.). One protein, PvLEA1, was located in the cell membrane in both cell lines (Fig. 1, Table 1). Two remaining proteins, PvLEA3 and PvLEA22, were distributed in the ER/Golgi in the Pv11 cells (Fig. 1, S2 Fig.) and in the ER/Golgi and cytoplasm, respectively, in Sf9 (Fig. 1, S2 Fig.). We verified the ER/Golgi localization of PvLEA3 and PvLEA22 proteins via staining by Cytopainter kit (Abcam) (S3 Fig.). The revealed localization of PvLEA proteins in Pv11 cells remained the same after anhydrobiosis-inducing treatment by trehalose for 24 hours.

**Table 1.**
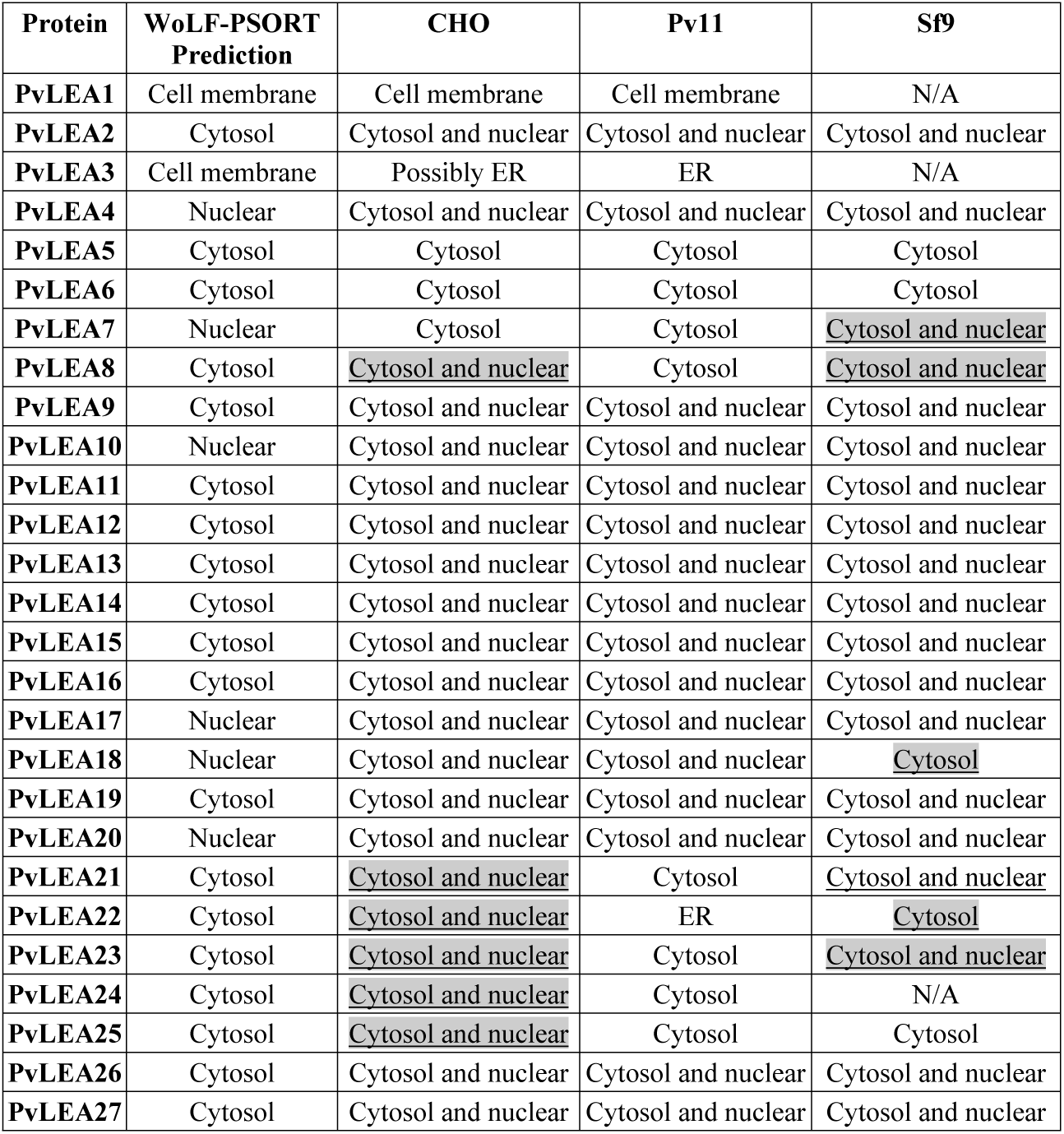
Comparison of PvLEA proteins’ subcellular localization in different cell cultures and WoLF PSORT predictions.We predicted the types of subcellular localization of different PvLEA proteins using WoLF PSORT and obtained e xperimental data with corresponding PvLEA(x)-AcGFP1 chimers in the CHO, Pv11, and Sf9 cells (x = 1-27). N/A – three PvLEA(x)-AcGFP chimers (with *PvLea1, PvLea3*, and *PvLea24* genes) caused aberrations in cell shape and toxicity in Sf9 cells. We marked revealed differences in localization in CHO or Sf9 cells in comparison to Pv11 cells by underscoring in the corresponding table cells and grey background color. Data on localization in CHO cells and WoLF PSORT predictions are in accordance with (7).

| Protein | WoLF-PSORT Prediction | CHO | Pv11 | Sf9 |
| --- | --- | --- | --- | --- |
| PvLEA1 | Cell membrane | Cell membrane | Cell membrane | N/A |
| PvLEA2 | Cytosol | Cytosol and nuclear | Cytosol and nuclear | Cytosol and nuclear |
| PvLEA3 | Cell membrane | Possibly ER | ER | N/A |
| PvLEA4 | Nuclear | Cytosol and nuclear | Cytosol and nuclear | Cytosol and nuclear |
| PvLEA5 | Cytosol | Cytosol | Cytosol | Cytosol |
| PvLEA6 | Cytosol | Cytosol | Cytosol | Cytosol |
| PvLEA7 | Nuclear | Cytosol | Cytosol | Cytosol and nuclear |
| PvLEA8 | Cytosol | Cytosol and nuclear | Cytosol | Cytosol and nuclear |
| PvLEA9 | Cytosol | Cytosol and nuclear | Cytosol and nuclear | Cytosol and nuclear |
| PvLEA10 | Nuclear | Cytosol and nuclear | Cytosol and nuclear | Cytosol and nuclear |
| PvLEA11 | Cytosol | Cytosol and nuclear | Cytosol and nuclear | Cytosol and nuclear |
| PvLEA12 | Cytosol | Cytosol and nuclear | Cytosol and nuclear | Cytosol and nuclear |
| PvLEA13 | Cytosol | Cytosol and nuclear | Cytosol and nuclear | Cytosol and nuclear |
| PvLEA14 | Cytosol | Cytosol and nuclear | Cytosol and nuclear | Cytosol and nuclear |
| PvLEA15 | Cytosol | Cytosol and nuclear | Cytosol and nuclear | Cytosol and nuclear |
| PvLEA16 | Cytosol | Cytosol and nuclear | Cytosol and nuclear | Cytosol and nuclear |
| PvLEA17 | Nuclear | Cytosol and nuclear | Cytosol and nuclear | Cytosol and nuclear |
| PvLEA18 | Nuclear | Cytosol and nuclear | Cytosol and nuclear | Cytosol |
| PvLEA19 | Cytosol | Cytosol and nuclear | Cytosol and nuclear | Cytosol and nuclear |
| PvLEA20 | Nuclear | Cytosol and nuclear | Cytosol and nuclear | Cytosol and nuclear |
| PvLEA21 | Cytosol | Cytosol and nuclear | Cytosol | Cytosol and nuclear |
| PvLEA22 | Cytosol | Cytosol and nuclear | ER | Cytosol |
| PvLEA23 | Cytosol | Cytosol and nuclear | Cytosol | Cytosol and nuclear |
| PvLEA24 | Cytosol | Cytosol and nuclear | Cytosol | N/A |
| PvLEA25 | Cytosol | Cytosol and nuclear | Cytosol | Cytosol |
| PvLEA26 | Cytosol | Cytosol and nuclear | Cytosol and nuclear | Cytosol and nuclear |
| PvLEA27 | Cytosol | Cytosol and nuclear | Cytosol and nuclear | Cytosol and nuclear |

**Fig. 3.**
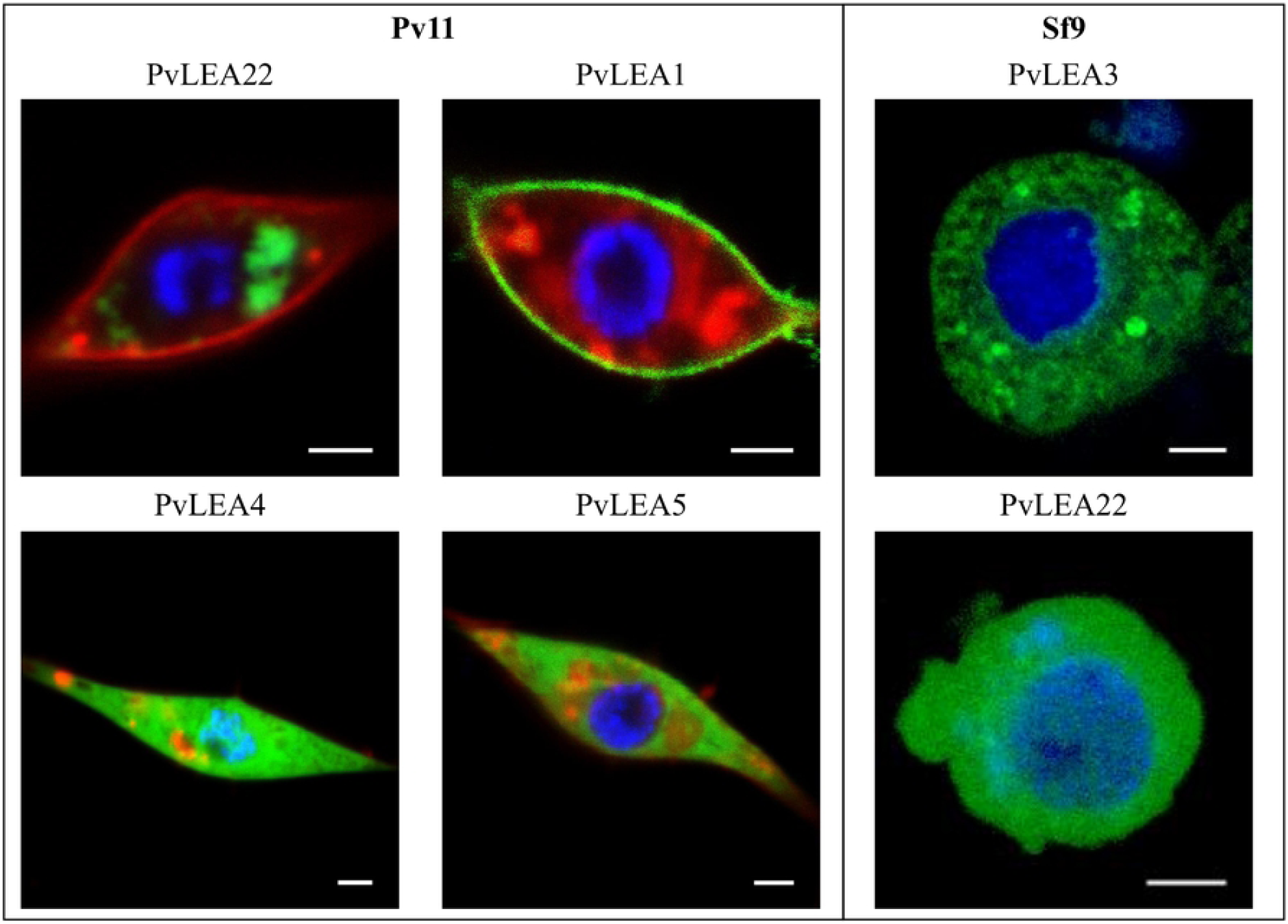
Representative images of PvLEA proteins localization in Pv11 and Sf9 cells. (Pv11) Localization of PvLEA(x)-AcGFP1 chimers in Pv11 cells: ER/Golgi (PvLEA22), membrane (PvLEA1), cytosol-nuclear (PvLEA4), and cytosol-only (PvLEA5). (Sf9) Localization of PvLEA(x)-AcGFP1 chimers in Sf9 cells: cytosol-nuclear (PvLEA8) and cytozol-only (PvLEA6). PvLEA proteins are indicated above image panels. ER/Golgi localization was verified using ER/Golgi staining (see methods).

Our current data on PvLEA protein localization in both Pv11 and Sf9 cells differ from our previously published data on CHO cells (7). The most widespread difference was the exclusion of five PvLEA proteins (PvLEA8, PvLEA21, PvLEA23, PvLEA24, and PvLEA25) from the nuclei of Pv11 cells compared to whole-cell distribution in CHO cells (Table 1). PvLEA22 in Pv11 cells was located in ER/Golgi in contrast to the whole cell (CHO cells) or cytosol only (Sf9). Localization of five other PvLEA proteins was also different between the Sf9 and Pv11 cells (Table 1).

## Discussion

Previously we showed that *PvLea* genes are expressed in *P. vanderplanki* larvae and become induced in response to desiccation (4). However, it was unknown whether anhydrobiotic phenotypes of these cells are also associated with an expression of the full range of PvLEA proteins. In this study, we analyzed *PvLea* gene expression in Pv11 cells. We found that all 27 previously identified *PvLea* genes were expressed in Pv11 cells except *PvLea16* and *PvLea17*. Similarly to larvae, anhydrobiosis onset in Pv11 cells is linked to induction of mRNA expression for the majority of *PvLea* genes. In Pv11 cells, we mediated anhydrobiosis through trehalose treatment for 48 hours, followed by a rapid desiccation (5). This procedure mimicked anhydrobiosis in *P. vanderplanki*’s larvae, which was successfully induced by slow desiccation taking nearly two days (16). During desiccation, larvae accumulate trehalose produced by an organ called the fat body and, at greatly elevated rates, express a multitude of genes encoding protective proteins. The fact that most *PvLea* genes achieve highest expression in Pv11 cells during trehalose treatment is well compatible with this model. Pv11 cells show considerable preservation of expression patterns of *PvLea* genes in comparison to larvae, both in a normal state and in anhydrobiosis. This preservation suggests that LEA proteins, which are highly expressed in whole larvae, are also necessary for successful anhydrobiosis in the Pv11 cells model.

One of the most expressed LEA-encoding genes in Pv11 cells during anhydrobiosis induction is *PvLea4* (Fig. 1). The corresponding PvLEA4 protein has already attracted research attention for its high level of expression of *PvLea* in *P. vanderplanki* larvae. It was shown to limit growth of aggregating protein particles, thereby acting as a molecular shield (17). This function is believed to be the main function of LEA proteins. Being typically disordered proteins, group 3 LEA proteins lack structure in normal conditions based on an abundance of hydrophilic amino acid residues (18). However, they obtain structure when water content decreases (19). The presence of several copies of a specific 11-mer motif is likely the main feature of LEA group 3 proteins, ensuring their molecular shield activity (20). Remarkably, in this study, we found that, in Pv11 cells, the presence of this motif (LEA_4 motif, Pfam ID: PF02987) in PvLEA proteins had some correlation with expression of the corresponding gene (S1 Table).

However, some PvLEA proteins lack the LEA_4 motif, raising the question of their functionality in *P. vanderplanki*’s anhydrobiosis (7). Observed difference in PvLEA proteins characteristics (7), together with the large difference in their expression and the fact that PvLEA proteins substantially diverged in sequence, suggest their functional specialization. Similarly to the case of *P. vanderplanki*, the presence of a multitude of LEA-encoding genes in some species is well established: for example, the genome of *Arabidopsis thaliana* encodes 51 different LEA proteins (21). The role of this diversity remains to be revealed, and one currently investigated aspect of LEA proteins’ specialization is their localization (22).

In this study, we expressed 27 PvLEA proteins from *P. vanderplanki* in fusion with AcGFP1 proteins in Pv11 and Sf9 cells. We visualized their localization using a confocal microscope, showing four different targeting types: cytosol, cytosol and nucleus, ER, and cytoplasmic membrane. We found differences in PvLEA preservation between Pv11 cells and Sf9 cells. For some proteins, localization was not preserved in these cell lines in comparison to CHO cells (7). Because Pv11 cells are derived from the host organism *P. vanderplanki*, localization of PvLEA proteins in this cell line is the most representative of their real targeting in *P. vanderplanki*. Found discrepancies in PvLEA proteins’ localization between different cell cultures raise important potential implications for the use of PvLEA proteins in dry preservation biotechnologies.

The function of proteins depends critically on their localization because chemical environments in different organelles differ. Thus, the protective (or any other) function of PvLEA proteins is also dependent on their correct localization. This can be important when a given protein is not reaching proper localization or is mislocalized. All revealed discrepancies between Pv11 and other cells are potential examples of protein mislocalization (Table 1), because localization in Pv11 cells was stricter than in other cells. Such protein mislocalization can be detrimental because it occurs in many cancers (23).

The main difference in PvLEA proteins’ localization between different cells in this work was the presence or absence of nuclear localization, which is a process governed by proteins, mostly karyopherin β family members referred to as importins (24). Thus, differences in PvLEA nuclear transport between different cells can be simply accounted by differences in structure of these nuclear-trafficking proteins. Differences in importins’ regulation and expression can be also a source of protein mislocalization (23).

Interestingly, in the Pv11 cells, none of the PvLEA proteins had mitochondrial targeting, despite mitochondria being potentially a source of increased production of reactive oxygen species during water loss based on the disruption of oxidative phosphorylation (25). In addition, the release of proteins from the intermembrane mitochondria space is a signal for the cells to undergo apoptosis (26). This reflects an increased demand for protection of these organelles in desiccating cells, which can explain the mitochondrial targeting of some LEA proteins (18).

## Conclusions

Our data provide a first glimpse into expression of *PvLea* in Pv11 cells. Pv11 cells express 25 out of 27 *P. vanderplanki* LEA-encoding genes, for which the expression pattern is similar to that in *P. vanderplanki* larvae. Out of these genes, 24 are upregulated in response to anhydrobiosis-inducing trehalose treatment. These data reinforce an association of *PvLea* genes identified in *P. vanderplanki* to its anhydrobiotic phenotype. We also showed that localization of PvLEA proteins in Pv11 cells is not always reproduced in other cell types. Such spatial aspects of protective proteins’ function should be taken into account when using PvLEA proteins in the emerging field of engineering of the artificial anhydrobiotic phenotype.

## Acknowledgments

We thank S. Kuznetsova and the Interdisciplinary Center for Analytical Microscopy of Kazan Federal University for the assistance in the PvLEA localization imaging. We are deeply grateful to Tomoe Shiratori (NARO) and Yuki Sato-Kikuzato (NARO) for their technical support.

## Supporting information captions

**S1 Fig. LEA genes’ expression in *P. vanderplanki* larvae and Pv11 cells in control and at different stages of anhydrobiosis induction.** Height of the bars depicts mean expression for replicates, whereas error bars show corresponding standard deviation. Colors of the bars indicate data for larvae (red) or Pv11 cells (blue-green). Genes are ordered in accordance with expression in larvae, and their names are indicated below the plot. Spearman correlation of expression means and the corresponding p-value are indicated by text on the plot. The plot is faceted in accordance with different experimental conditions, as indicated on the top of each panel. In the case of larvae, anhydrobiosis induction is represented by slow desiccation, which takes nearly 48 hours up to an air-dry state. In the case of Pv11 cells, which are unable to synthesize trehalose, anhydrobiosis induction consists of trehalose treatment for 48 hours, followed by a rapid desiccation (see Methods).

**S2 Fig. Plot of *PvLea* gene expression in Pv11 cells versus GRAVY index of hydropathy or number of LEA_4 motifs in respective protein.** The GRAVY index is the grand average of hydropathy index, describing proteins solubility. Negative GRAVY index values indicate hydrophilic characteristics of the corresponding protein. Used GRAVY index data and number of LEA_4 motifs (ID PF02987 in Pfam 26.0 database) are from (7). Mean values of expression for each gene are on x-axis.

**S1 Table. Correlation of PvLEA proteins characteristics with expression of corresponding genes in Pv11 cells.** Table is sorted in accordance with the correlation value. Rs: values of Spearman correlation. LEA_4 motifs: ID PF02987 in Pfam 26.0 database. GRAVY index - the grand average of hydropathy index, describing protein solubility. Negative values of the GRAVY index indicate hydrophilic characteristics of the corresponding protein. Used data on number of LEA_4 motifs and the GRAVY index come from (7).

**S3 Fig. Images of subcellular localization of PvLEA(x)-AcGFP1 chimers in Pv11 and Sf9 cells**

**S4 Fig. ER/Golgi staining of Pv11 cells expressing PvLEA3-AcGFP1 and PvLEA22-AcGFP1 chimers.** Pv11 cells expressing PvLEA3 and PvLEA22 with endoplasmic reticulum (ER) and Golgi apparatus stained by the CytoPainter Staining Kit (Abcam, USA) and DNA stained with Hoechst 33258 (Sigma, USA).

